# The Chemical Features of Polyanions Modulate Tau Aggregation and Conformational States

**DOI:** 10.1101/2022.07.28.501920

**Authors:** Kelly M. Montgomery, Emma C. Carroll, Aye Thwin, Paige Hodges, Daniel R. Southworth, Jason E. Gestwicki

**Affiliations:** Department of Pharmaceutical Chemistry, University of California San Francisco, San Francisco, CA 94158; The Institute for Neurodegenerative Diseases, University of California San Francisco, San Francisco, CA 94158; Department of Biochemistry and Biophysics, University of California San Francisco, San Francisco, CA 94158

## Abstract

The aggregation of tau into insoluble fibrils is a defining feature of neurodegenerative tauopathies. However, tau has a positive overall charge and is highly soluble; so polyanions, such as heparin, are typically required to promote its aggregation *in vitro*. There are dozens of polyanions in living systems and it is not clear which ones might promote this process. Here, we systematically measure the ability of 30 diverse, anionic biomolecules to initiate tau aggregation, using either wild type (WT) tau or the disease associated P301S mutant. We find that polyanions from many different structural classes can promote fibril formation and that P301S tau is sensitive to a greater number of polyanions (19/30) than WT tau (16/30). We also find that some polyanions preferentially reduce the lag time of the aggregation reactions, while others enhance the elongation rate, suggesting that they act on partially distinct steps. From the resulting structure-activity relationships, the valency of the polyanion seems to be an important chemical feature, such that anions with low valency tend to be weaker aggregation inducers, even at the same overall charge. Finally, the identity of the polyanion influences fibril morphology, based on electron microscopy and limited proteolysis. These results provide insight into the crucial role of polyanion—tau interactions in modulating tau conformational dynamics with implications for understanding the tau aggregation landscape in a complex cellular environment.

## Introduction

The class of neurodegenerative disorders known as tauopathies, including Alzheimer’s disease (AD), cortical basal degeneration (CBD) and progressive supranuclear palsy (PSP), are characterized by the accumulation of insoluble protein aggregates in the brain^1,2^. These aggregates are primarily composed of microtubule-associated protein tau (MAPT/tau), an intrinsically disordered protein that is expressed as a series of six distinct splice isoforms^3,4^. Tau’s isoforms are composed of a variable number of N-terminal domains (0N, 1N or 2N), a proline-rich domain and either three or four microtubule-binding repeats (3R or 4R; Fig 1A). The common adult isoform of tau, 0N4R, is strongly cationic at physiological pH, with an isoelectric point of ∼9.5. Accordingly, it has been known for decades that purified tau is highly soluble and not prone to spontaneously self-associate *in vitro*, even at extremes of pH and temperature^5^. Rather, tau aggregation is typically initiated by the addition of polyanionic biomolecules, such as heparin sodium (HS), which leads to relatively rapid self-assembly^6^. It is thought that the polyanions reduce charge repulsion between cationic tau monomers, allowing the juxtaposition of aggregation motifs within the R repeats^7,8^. These observations suggest that polyanions could also be involved in the initiation of tau aggregation *in vivo*. In support of this idea, HS is associated with tau pathology in patient brains^9^ and unresolved densities are observed in some purified, patient-derived tau fibril samples, which are hypothesized to be anions or salts^10,11^. At present, we do not know the identity of the critical natural anion(s) or how their chemical properties might contribute to tau aggregation or fibril structure.

**Figure 1.**
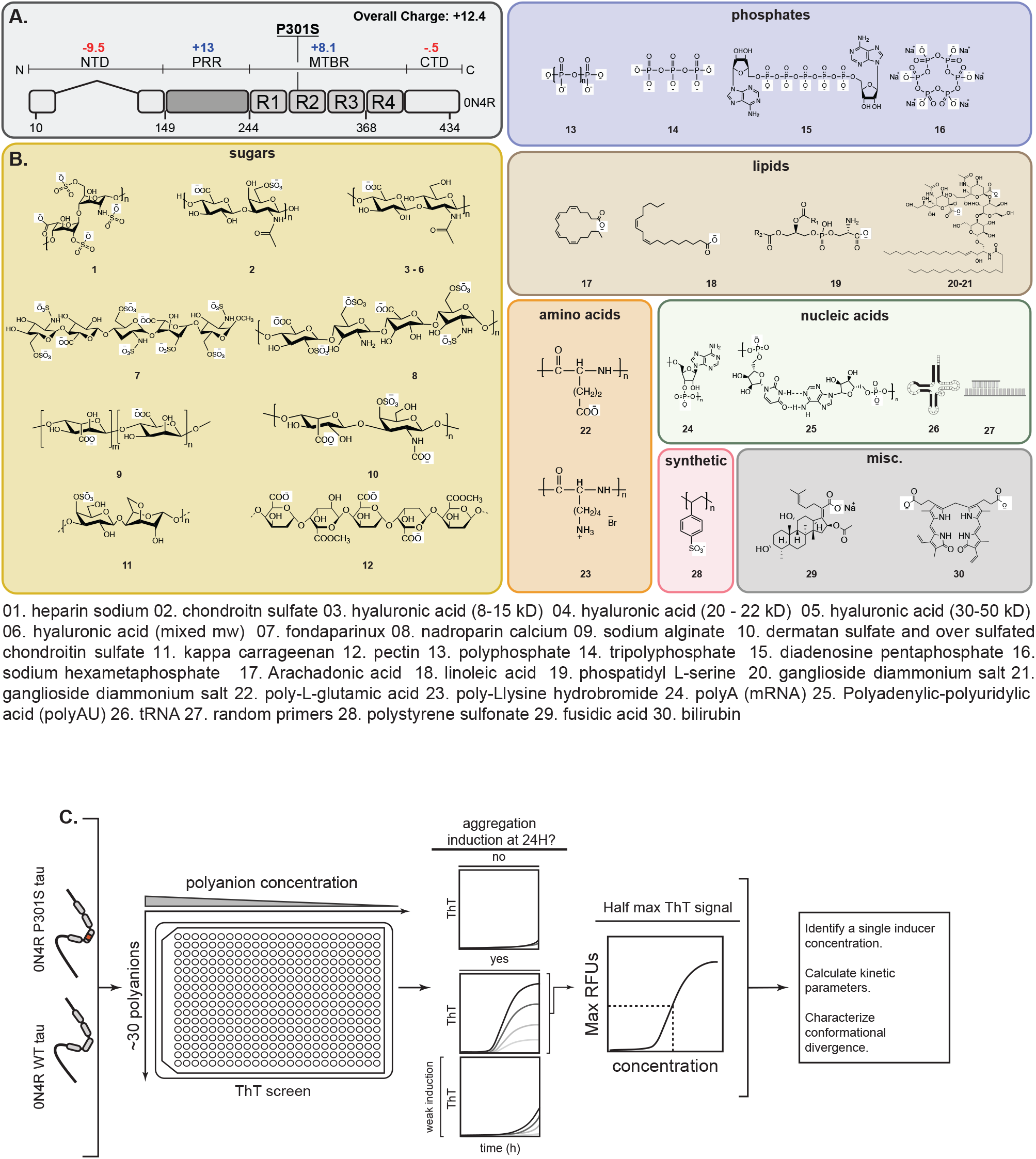
Workflow for testing the effects of a polyanion library on tau self-assembly. (A) The domain architecture of the longest adult isoform of tau (0N4R). The overall charge of the domains is indicated and the location of the P310S missense mutation is shown. N-terminal domain (NTD); proline-rich region (PRR); microtubule-binding repeats (MTBR); C-terminal domain (CTD). (B) Chemical structures of 30 anionic biomolecules, grouped by series. When appropriate, the minimal repeating unit is shown, and the average polymer length (n) is indicated in the Supporting Information. Compounds 26 and 27 are shown as cartoons because they do have repeating structure (see Methods). (C) The workflow for screening the anion library. Briefly, 0N4R tau (WT) and 0N4R P301S mutant tau (P301S) proteins were first tested against a range of concentrations of each library member in ThT assays. Anions were excluded from further analysis if they produced artifacts (Supplemental Fig. 1) or if they failed to produce a ThT signal 40 RFUs above baseline fluorescence (RFU ≤ 40) at 24 hours. For the remaining molecules, the half-maximal effective concentration (NC50) was determined, and subsequent kinetic studies performed at that anion concentration. From those studies, the kinetic parameters, including lag-time and elongation rate were determined.

Structural studies have revealed that the core of tau can adopt a variety of conformations within fibrils; for instance, postmortem brain slices derived from patients with PSP and CBD contain tau fibrils with distinct folds^12–15^. Could the identity of the anion help dictate these specific conformations? To reach the fibril state, tau is known to transition through intermediate structures, including oligomers^16,17^. While many factors likely contribute to the eventual structural differences between fibrils^18–20^, we hypothesize that the polyanion could help guide early stages of the self-assembly process and contribute to determining the final structure. Indeed, an increasing body of evidence suggests that polyanions have an effect on early stages of fibril nucleation *in vitro*^21–23^. Yet, only a limited number of anions, including those in the broad categories of sugars^8^, fatty acids^24^, nucleic acids^25^, and phosphates^26,27^ have been studied for their ability to promote tau self-assembly and a systematic study, involving direct comparisons between these varied molecules under the same experimental conditions, is lacking.

Here, we collected 30 chemically and structurally diverse anions and tested them side-by-side in a thioflavin T (ThT) assay to identify those that mediate tau’s aggregation. We find that a surprisingly large number of anionic biomolecules, including certain naturally occurring sugars, polypeptides, nucleic acids, amino acids and lipids, promote this process. Valency appears to be an important feature of these molecules, because only polyanions of sufficient repeat length were able to promote fibril formation. We also find that a disease-associated mutant, P301S tau, is sensitive to a larger number of anions (19/30) than wild type (16/30), which could be one reason why it is linked to severe disease. To explore the potential impact of polyanions on the conformation of tau fibrils, we selected some of the most potent inducers and explored the resulting fibrils by limited proteolysis and transmission electron microscopy (TEM). Remarkably, we find that the identity of the polyanion has a dramatic effect on tau fibril conformation. Together, these results expand our knowledge of the role of polyanions in tau aggregation *in vitro*. Based on these findings, we speculate that the availability of specific anions in the brain is one important factor in shaping fibril conformation.

## Results

### Creation of an anion library

In choosing anions for a chemical library, we sought to incorporate benchmark compounds, such as heparin sodium (HS), as well as molecules from a variety of structural classes that had never been previously tested for their effects on tau aggregation. Accordingly, we purchased anionic sugars (**1**-**12**), polyphosphates (**13**-**16**), short chain fatty acids (**17**-**21**), polypeptides (**22**-**23**) and oligonucleotides (**24**-**27**; Fig 1B), as well as anions from more chemically diverse classes, such as antibiotics (**29**), biladienes (**30**) and synthetic polymers (*i*.*e* polystyrene sulfonate, **28**). Only a subset of these compounds (**1, 6, 13, 14**, and **24**), had, to our knowledge, been previously tested for their effects on tau aggregation. While the major goal of this panel was to sample different scaffolds, some of the library members varied in their chemical properties. For example, fondaparinux (**7**), polyphosphate (**13**), and polyadenylic-polyuridylic acid (poly-AU; **25**) have varying charge density: -10 per monomer for compound **7**, -1 per monomer for **13** and -2 for **25** (Fig 1B). Thus, we also hoped that screening this collection might also begin to reveal chemical features important for tau aggregation.

Previous work had shown that anions employ at least two mechanisms to promote tau self-assembly. For example, HS (**1**) is an example of the polymer class of inducers and it is decorated with sulfates from a repeating disaccharide unit (Fig 1B). HS and related compounds are thought to mitigate the unfavorable, long-range electrostatic interactions in tau, favoring hydrophobic collapse of the core^5,28–30^. However, another mechanism is linked to monovalent anions, such as arachidonic acid (**17**), which are thought to function by first forming micelles on which tau assembles^31–33^. In our anion library, examples of both categories are included, and we anticipated that polymers, such as sugars, nucleotides, and phosphates, might function similarly to HS, while the lipids and other non-polymers, might function as micelles.

### Screens identify the subset of anions that promote tau aggregation

To evaluate the anion library, we measured tau aggregation using a 384-well, plate-based, ThT platform (Fig 1C).^24^ Briefly, our goal was to first screen each anion to reveal which ones could support tau aggregation; then, we would focus on the most potent inducers to perform more detailed kinetic and structural studies. In these experiments, we employed two purified, recombinant human proteins; 0N4R tau (WT; Table 1) and a 0N4R P301S mutant (P301S; Table 2). P301S is a genetic mutation associated with FTD-linked Parkinsonism-17 (FTDP-17) that produces severe frontal temporal atrophy.^34^ It is known that P301S is more aggregation prone than WT in the presence of HS^35,36^ so we reasoned that it might be more sensitive to weaker inducers.

**Table 1.**
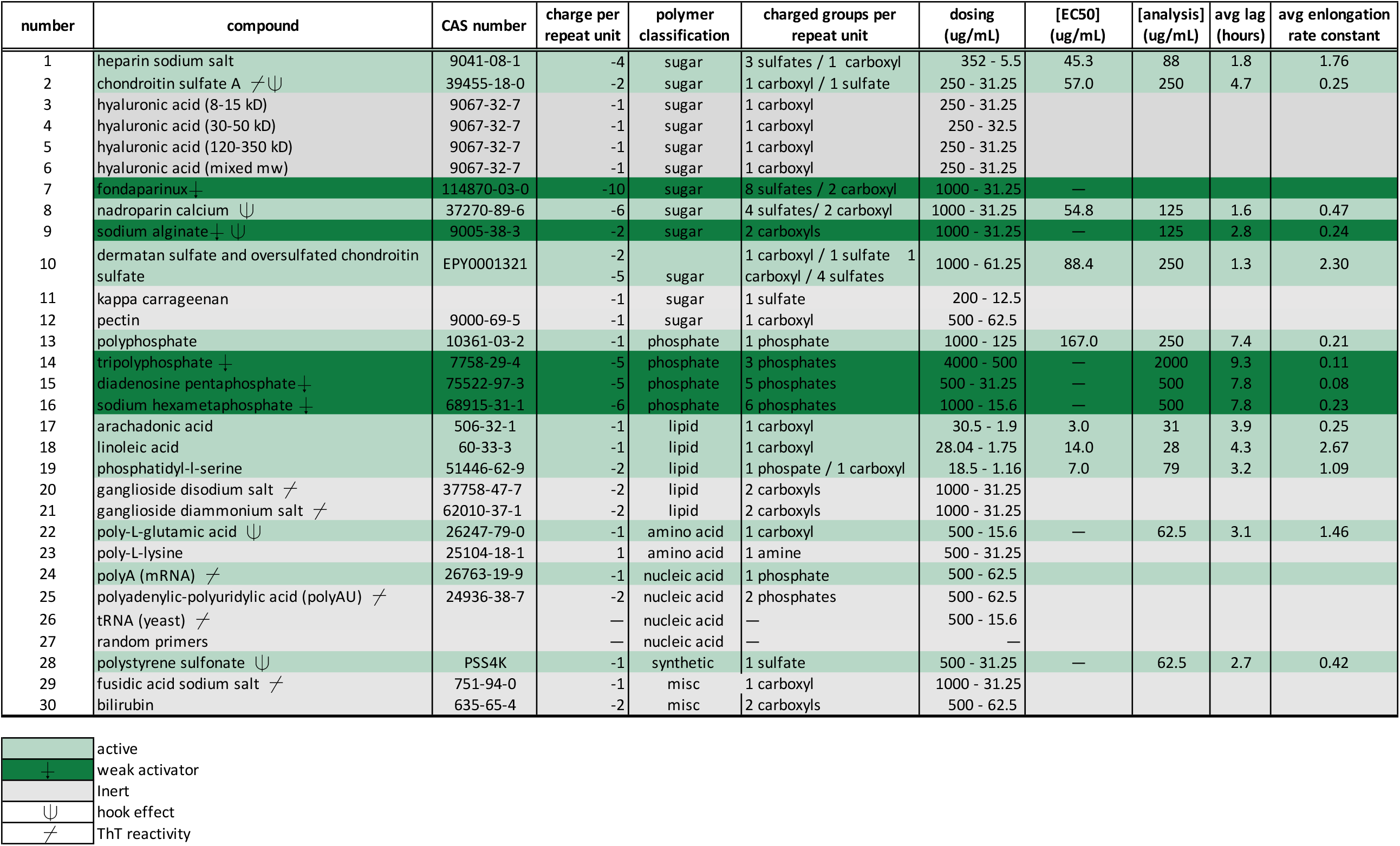
Effects of anions on WT tau aggregation. For each anion, a compound number and CAS number are included, along with a description of its chemical class (*e*.*g*., sugar), type of negative charge group and number of charges per repeat. The screening concentrations that were tested are also shown (dosing, µg/mL). Anions that produce ThT positive species (?RFU > 40), are considered active (green) and inactive ones are those that produce (?RFU ≤ 40) (grey). A subset of inducers, denoted by (⌿), promote aberrant ThT fluorescence in the absence of protein (Supplemental Fig 1). A subset of active anions produce only weak aggregation and are denoted by (⍖). Molecules that produce hook-like affects (Supplemental Fig 4) are denoted by (⍦).

**Table 2.**
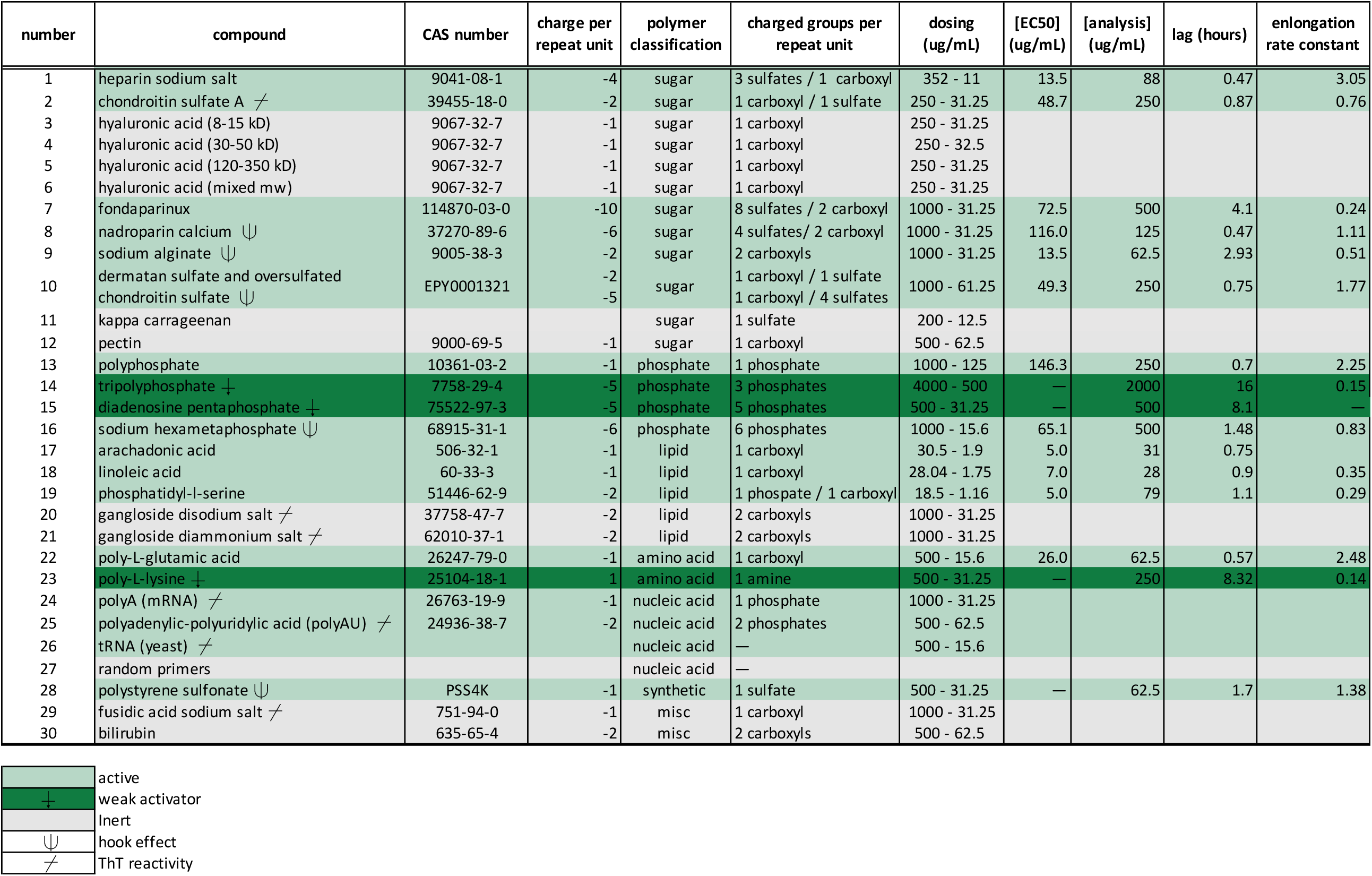
Effects of anions on P301L tau aggregation. For each anion, a compound number and CAS number are included, along with a description of its chemical class (*e*.*g*., sugar), type of negative charge group and number of charges per repeat. The screening concentrations that were tested are also shown (dosing, µg/mL). Anions that produce ThT positive species (?RFU > 40), are considered active (green) and inactive ones are those that produce (?RFU ≤ 40) (grey). A subset of inducers, denoted by (⌿), promote aberrant ThT fluorescence in the absence of protein (Supplemental Fig 1). A subset of active anions produces only weak aggregation and are denoted by (⍖). Molecules that produce hook-like affects (Supplemental Fig 4) are denoted by (⍦).

| number | compound | CAS number | charge per repeat unit | polymer classification | charged groups per repeat unit | dosing (ug/mL) | [EC50] (ug/mL) | [analysis] (ug/mL) | lag (hours) | enlongation rate constant |
| --- | --- | --- | --- | --- | --- | --- | --- | --- | --- | --- |
| 1 | heparin sodium salt | 9041-08-1 | -4 | sugar | 3 sulfates / 1 carboxyl | 352 - 11 | 13.5 | 88 | 0.47 | 3.05 |
| 2 | chondroitin sulfate A $\nrightarrow$ | 39455-18-0 | -2 | sugar | 1 carboxyl / 1 sulfate | 250 - 31.25 | 48.7 | 250 | 0.87 | 0.76 |
| 3 | hyaluronic acid (8-15 kD) | 9067-32-7 | -1 | sugar | 1 carboxyl | 250 - 31.25 |  |  |  |  |
| 4 | hyaluronic acid (30-50 kD) | 9067-32-7 | -1 | sugar | 1 carboxyl | 250 - 32.5 |  |  |  |  |
| 5 | hyaluronic acid (120-350 kD) | 9067-32-7 | -1 | sugar | 1 carboxyl | 250 - 31.25 |  |  |  |  |
| 6 | hyaluronic acid (mixed mw) | 9067-32-7 | -1 | sugar | 1 carboxyl | 250 - 31.25 |  |  |  |  |
| 7 | fondaparinux | 114870-03-0 | -10 | sugar | 8 sulfates / 2 carboxyl | 1000 - 31.25 | 72.5 | 500 | 4.1 | 0.24 |
| 8 | nadroparin calcium $\Psi$ | 37270-89-6 | -6 | sugar | 4 sulfates/ 2 carboxyl | 1000 - 31.25 | 116.0 | 125 | 0.47 | 1.11 |
| 9 | sodium alginate $\Psi$ | 9005-38-3 | -2 | sugar | 2 carboxyls | 1000 - 31.25 | 13.5 | 62.5 | 2.93 | 0.51 |
| 10 | dermatan sulfate and oversulfated chondroitin sulfate $\Psi$ | EPY0001321 | -2<br>-5 | sugar | 1 carboxyl / 1 sulfate<br>1 carboxyl / 4 sulfates | 1000 - 61.25 | 49.3 | 250 | 0.75 | 1.77 |
| 11 | kappa carrageenan |  |  | sugar | 1 sulfate | 200 - 12.5 |  |  |  |  |
| 12 | pectin | 9000-69-5 | -1 | sugar | 1 carboxyl | 500 - 62.5 |  |  |  |  |
| 13 | polyphosphate | 10361-03-2 | -1 | phosphate | 1 phosphate | 1000 - 125 | 146.3 | 250 | 0.7 | 2.25 |
| 14 | tripolyphosphate $\vdash$ | 7758-29-4 | -5 | phosphate | 3 phosphates | 4000 - 500 | — | 2000 | 16 | 0.15 |
| 15 | diadenosine pentaphosphate $\vdash$ | 75522-97-3 | -5 | phosphate | 5 phosphates | 500 - 31.25 | — | 500 | 8.1 | — |
| 16 | sodium hexametaphosphate $\Psi$ | 68915-31-1 | -6 | phosphate | 6 phosphates | 1000 - 15.6 | 65.1 | 500 | 1.48 | 0.83 |
| 17 | arachadonic acid | 506-32-1 | -1 | lipid | 1 carboxyl | 30.5 - 1.9 | 5.0 | 31 | 0.75 |  |
| 18 | linoleic acid | 60-33-3 | -1 | lipid | 1 carboxyl | 28.04 - 1.75 | 7.0 | 28 | 0.9 | 0.35 |
| 19 | phosphatidyl-l-serine | 51446-62-9 | -2 | lipid | 1 phospate / 1 carboxyl | 18.5 - 1.16 | 5.0 | 79 | 1.1 | 0.29 |
| 20 | gangloside disodium salt $\nrightarrow$ | 37758-47-7 | -2 | lipid | 2 carboxyls | 1000 - 31.25 | | | | |
| 21 | gangloside diammonium salt $\nrightarrow$ | 62010-37-1 | -2 | lipid | 2 carboxyls | 1000 - 31.25 | | | | |
| 22 | poly-L-glutamic acid | 26247-79-0 | -1 | amino acid | 1 carboxyl | 500 - 15.6 | 26.0 | 62.5 | 0.57 | 2.48 |
| 23 | poly-L-lysine $\vdash$ | 25104-18-1 | 1 | amino acid | 1 amine | 500 - 31.25 | — | 250 | 8.32 | 0.14 |
| 24 | polyA (mRNA) $\nrightarrow$ | 26763-19-9 | -1 | nucleic acid | 1 phosphate | 1000 - 31.25 | | | | |
| 25 | polyadenylic-polyuridylic acid (polyAU) $\nrightarrow$ | 24936-38-7 | -2 | nucleic acid | 2 phosphates | 500 - 62.5 | | | | |
| 26 | tRNA (yeast) $\nrightarrow$ | | | nucleic acid | — | 500 - 15.6 | | | | |
| 27 | random primers |  |  | nucleic acid | — |  |  |  |  |  |
| 28 | polystyrene sulfonate $\Psi$ | PSS4K | -1 | synthetic | 1 sulfate | 500 - 31.25 | — | 62.5 | 1.7 | 1.38 |
| 29 | fusidic acid sodium salt $\nrightarrow$ | 751-94-0 | -1 | misc | 1 carboxyl | 1000 - 31.25 | | | | |
| 30 | bilirubin | 635-65-4 | -2 | misc | 2 carboxyls | 500 - 62.5 |  |  |  |  |

|  |  |
| --- | --- |
|  | active |
| $\vdash$ | weak activator |
|  | Inert |
| $\Psi$ | hook effect |
| $\nrightarrow$ | ThT reactivity |

In the initial screens, we tested each member of the anion library at a range of concentrations. These ranges (see Tables 1 and 2) were either selected from the literature (for known inducers) or chosen empirically (for those that had not been studied previously). These experiments were performed in triplicate using ThT signal measured every 5 minutes for a minimum of 24 hours with shaking at 37 °C (see Methods). At the same time, we performed experiments in the absence of tau protein to reveal any anions that might produce artifacts. Indeed, this control was important because we found that, for WT tau, gangliosides (**20-21**), fusidic acid (**29**) and several nucleic acids, including tRNA (**26**), polyA (**24**), and polyAU (**25**), produced ThT signal in the absence of protein (Supplemental Fig 1) and, accordingly, they were excluded from further analysis (Table 1). Chondroitin sulfate A (CS; **2**) produced a relatively modest tau-independent signal which could be subtracted from the experimental samples (Supplemental Fig 1), so this inducer was carried forward into the next experiments. For the remaining anions, we placed them into three categories based on the maximum ThT signal that they produced. Those considered to be “inactive” failed to reach saturation and yielded 40 or less RFUs (ΔRFU ≤ 40) above baseline at 24 hours (Tables 1 and 2; Supplemental Fig 2). Inducers were determined to be “moderately active” if they yielded a ΔRFU > 40, but did not reach a maximum ThT signal at the highest tested concentration of the anion (Supplemental Fig 3). Finally, inducers were “active” if they produced ΔRFU > 40 with a full sigmoidal curve. For these anions, we determined their relative potency by fitting the sigmoid of the dose response and determining the half-maximal value (EC_50_). Importantly, we noted that a subset of “active” anions produced a hook-like effect in their dose response; they initially promote ThT signal at lower concentrations and then become inhibitory with increased dosing (Supplemental Fig 4). For example, polystyrene sulfonate (**28**) becomes inhibitory at concentrations above 250 µg/mL. For this subset of molecules, we estimated EC_50_ using the upper inflection point as the top concentration.

**Figure 2.**
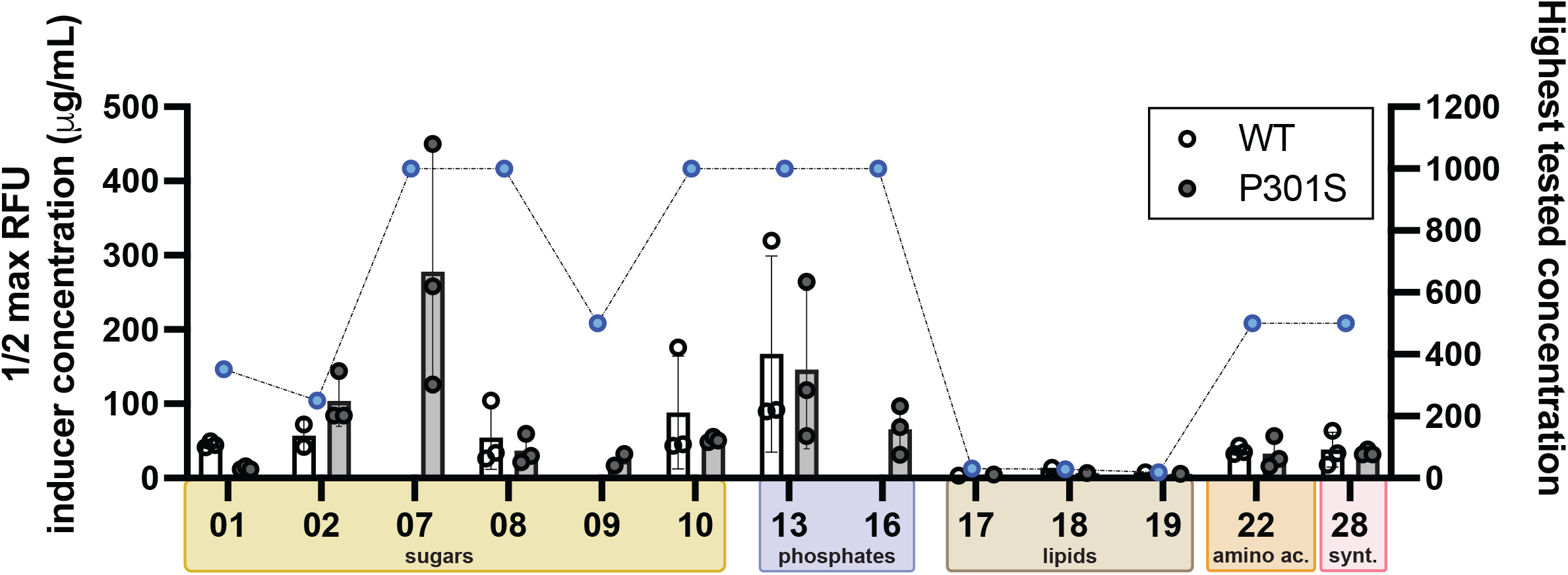
A large number of diverse anions can support tau fibril formation. (A) The potency values (half-maximal concentration; EC50) of the active anions are shown. To calculate potency, the highest RFU for each inducer concentration was extracted, plotted, and subsequently fit to a 4-parameter nonlinear regression in GraphPad Prism. Reported half-maximal concentration is a mean of EC50 values calculated from three independent experiments each conducted in technical triplicates (n=9) with error reported as SEM. (B) The insert shows the highest ThT fluorescence signal for each active inducer. Values are the average of three independent experiments performed in triplicate each (n=9) and error bars are SEM. For clarity, the results from the inactive and moderately active anions are not shown (see Tables 1 and 2).

**Figure 3.**
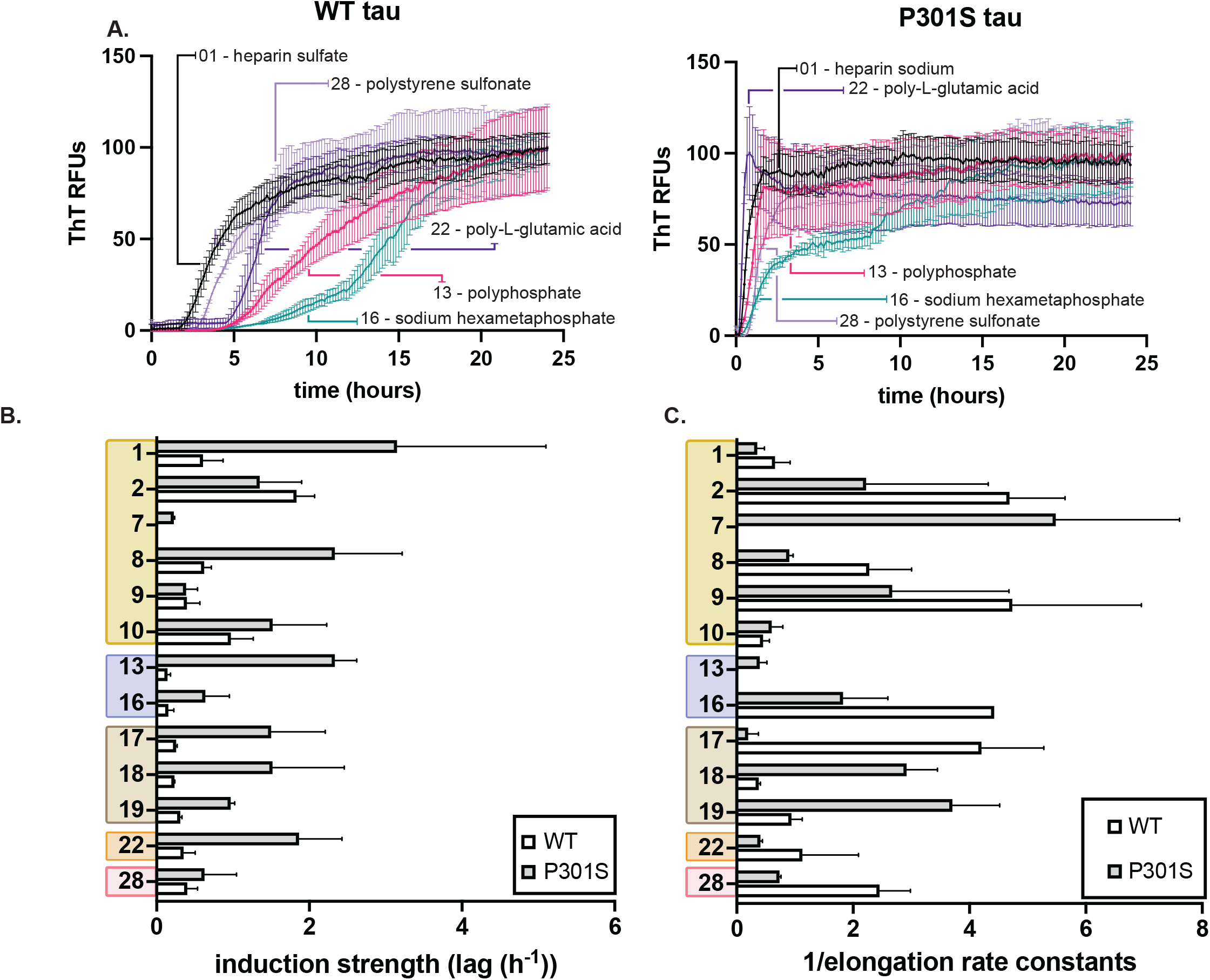
Anions have differential effects on lag time and elongation rate. (A) Representative ThT assay results, comparing heparin sodium (01), polyphosphate (13), sodium hexametaphosphate (15), poly-L glutamic acid (22), and polystyrene sulfonate (28) on recombinant 0N4R TauWT (left) and 0N4R Tau P301S (right) aggregation kinetics. The anions were used at their half-maximum concentration (see Tables 1 and 2) and tau proteins at 10 µM. Results are the average of at least three independent experiments performed in triplicate and the error bars represent SEM. For each result, the signal from control experiments using no tau was subtracted. (B) Anions have differential effects on lag time. Values are plotted as reciprocal (lag-1), termed the induction strength. Inactive inducers and those with weak signal were omitted from the analysis (see text). Results are the average of at least three independent experiments performed in triplicate and the error bars represent SEM. (C) From the same aggregation reactions, the elongation rate was calculated and plotted as the reciprocal (elongation rate-1), termed the elongation speed. Results are the average of at least three independent experiments performed in triplicate and the error bars represent SEM.

**Figure 4.**
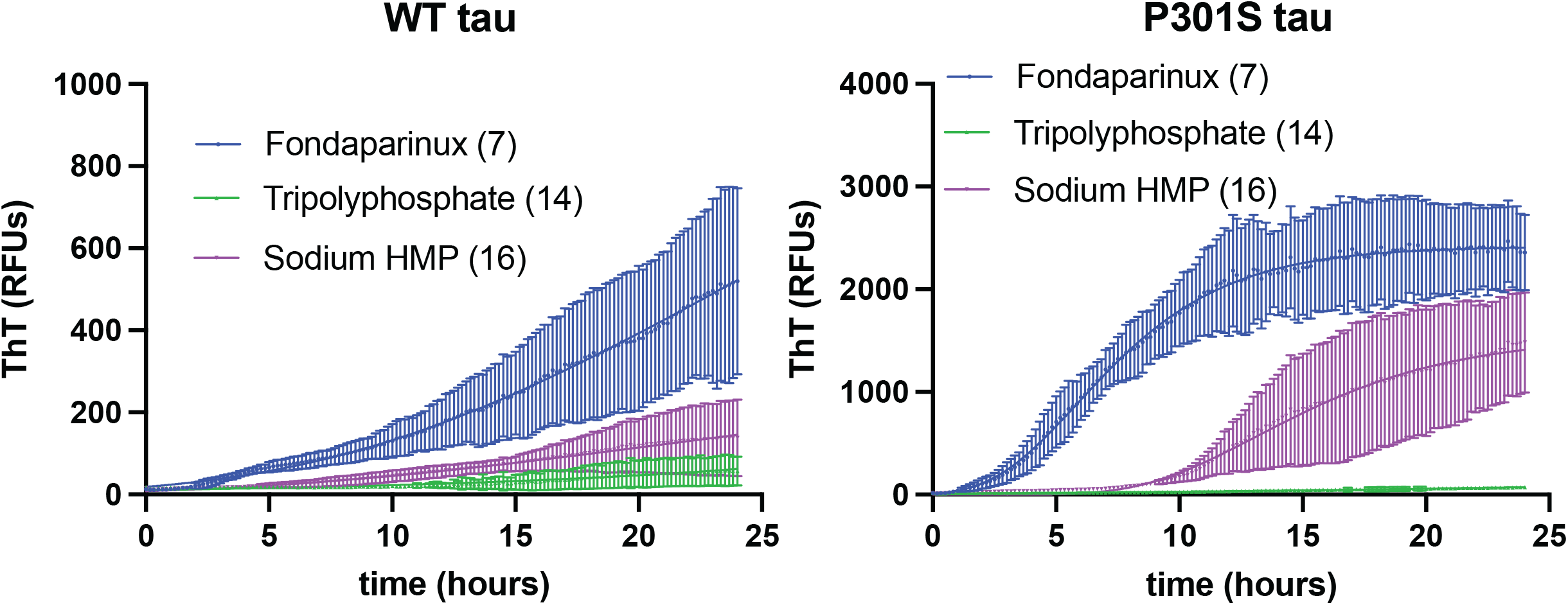
Polyanion valency is an important parameter in dictating tau fibril formation. 2N solutions of fondaparinux (7), tripolyphosphate (14) or sodium hexametaphosphate (16) were used to induce (left) WT and (right) P301S tau (10 µM) aggregation for 24 hours at 37 °C with constant shaking, and the resulting ThT curves were plotted. Results are the average of at least three experiments performed in triplicate and the error bars represent SEM (n=9).

One of the striking results from this screen is that tau aggregation is achieved with a large number of diverse anions. More specifically, for WT tau, we find that (11/30) molecules are strongly active and (5/30) are modestly active (Table 1). Together, these active molecules represent 6 out of the 7 structural categories (Fig 2), suggesting that many classes of anions, with a variety of backbones, can support tau self-assembly. However, these molecules were not all equally potent. Of the molecules tested, the sugars (**1, 2, 7-10**) and lipids (**17-19**) are generally the lowest EC_50_ values (*e*.*g*. active at the lowest concentrations; Fig 2), while phosphates (**13-16**) tend to be the least potent (*e*.*g*. require the highest concentrations). It is important to note that these comparisons are imperfect because the natural polymers used here are heterogeneous in length and valency, so it is difficult to compare their molar concentrations. Finally, we were surprised to find that several of the anions are “inert” (unable to produce ThT signal), including hyaluronic acids (**3-6**), kappa carrageenan (**11**), pectin (**12**), gangliosides (**20-21**), fusidic acid (**29**) and bilirubin (**30**) (Tables 1 and 2). It is not yet clear why these anions are inactive, but it is possible that they only produce amorphous aggregates that are not detectable by ThT. Regardless, this finding is worth highlighting because it shows that charge alone is not sufficient to drive the amyloid process.

In addition to having different potency values, we also find that anions produced different levels of maximal ThT signal (Table 1 and 2). Maximum ThT fluorescence is likely a product of the number of fibrils, the number of ThT binding sites in those structures and the chemical features of the binding sites (*e*.*g*. hydrophobicity).^37^ Thus, these results begin to suggest that the conformation(s) of the fibrils formed by the different anions might be distinct.

### Kinetic studies reveal the differential effects of anions on lag time and/or elongation rate

In aggregation reactions, the lag time is used to estimate the time required for an inducer to initiate formation of small oligomers, while the elongation rate is representative of multiple steps, including monomer addition and fibril fragmentation.^38^ Thus, we reasoned that comparing these values for reactions initiated by different anions might provide further insight into the steps that are affected. In these experiments, we used each anion at its EC_50_ concentration, using triplicate wells and repeating the studies three times with independent tau protein samples (n=9, three biological replicates). From the resulting data, the lag time and elongation rate were determined with the Gompertz function using the grace plotting program (see Methods). To facilitate comparisons between anions, we plotted the reciprocal of the lag time (lag time^-1^) to calculate an “induction strength”; where higher values are indicative of faster aggregation (Fig 3). Although there is considerable variability within classes, we find that sugars generally initiate fibril formation the fastest, with lag times around 2 to 2.5 hours for WT tau (Table 1). In contrast, the polyphosphates promote aggregation with considerably longer lag times (∼4 to 16 hours). Next, we similarly plotted the reciprocal of the elongation rate for each reaction; where high values are indicative of faster progression of fibril assembly. Again, there are differences between and within chemical classes, but sugars tend to produce the fastest elongation speeds. With both sets of values in-hand, we could then identify anions that might preferentially impact lag time or elongation rate. Indeed, we noted that HS (**1**) tended to produce dramatic effects on lag time, with comparatively little effect on elongation rate, as previously reported.^7^ However, other sugars, such as **2** and **8**, had relatively strong effects on both steps. Other compounds, such as **7** and **9**, had a disproportionate impact on elongation rate. Thus, the identity of an anion seems to determine which steps in the aggregation process are most impacted.

### Mutant P301S is sensitive to a wider range of anions

Although we have, to this point, focused on the results obtained with WT tau, these experiments were also performed, in parallel, using mutant P301S tau. A comparison between the results obtained with these two proteins suggested that, as expected, most anions that induce weak or moderate ThT signals for WT tau give relatively stronger signals using the P301S construct, with shorter lag times (Table 2). For example, fondaparinux (7) was a weak/modest activator for WT tau, but a strong one for P301S (Tables 1 and 2). Moreover, for a few anions, such as **25**, aggregation was only detected with P301S. We also noted that even the cation, poly-L-lysine, which was originally selected as a negative control, was able to weakly stimulate ThT signal for P301S. Crowding agents are known to accelerate aggregation^39^, so it is possible that poly-L-lysine might operate by this mechanism. Another unexpected result was that a small number of anions, such as **17** and **28**, produced relatively faster elongation rates for WT vs. P301S (Fig 3). Regardless, in most cases, we found that P301S was more sensitive than WT for nearly all of the anions. These findings support a model in which the strong link between this mutation and FTD could be, in part, a product of its sensitivity to a wider range of naturally occurring anions.

### Anions require a combination of both charge density and valency to support tau aggregation

In these screens, we found that molecules from a surprising number of chemical classes could support tau assembly. Therefore, it seems likely that the individual details of the scaffold backbone (*e*.*g*. sugar, peptide, *etc*.) might be relatively less important than their shared physical features, such as charge and valency. Indeed, careful studies using purified heparins and polyphosphates have also pointed to a key role for valency in tau aggregation^26^. To test this idea in more detail, we obtained additional anions, including monomeric ones, within the sugar and phosphate chemical series and tested them in ThT assays. Within these series of chemically defined molecules, we compared their activity on a per monomer, molarity basis. The results supported the idea that anion valency seems to be an essential feature of inducers. For example, tripolyphosphate **(14)** is nearly inactive compared to the penta- and hexavalent compounds: fondaparinux **(7)** and sodium hexametaphosphate **(16)**, respectively (Fig 4), even at the same monomer molarity. As previously suggested,^7^ we speculate that multivalent anions might bridge multiple tau monomers, increasing their local concentration and, ultimately, enhancing self-assembly. However, valency is clearly not the only important feature because highly valent compounds with sparse charge, such as hyaluronic acids (**3-6**), are ineffective inducers, suggesting that an optimal balance of charge density and polymer valency may be important.

### The identity of the polyanion dictates fibril structure

In addition to their effects on kinetics, we hypothesized that the identity of the polyanion might impact fibril structure. To ask this question, we turned to limited proteolysis. The advantage of this approach is that it reveals potential differences in the “fuzzy coat” of tau fibrils (*e*.*g*. regions outside of the well-folded core), a region that makes up the bulk of tau fibrils and has been shown to adopt heterogeneous conformers^14,40–43^. Accordingly, WT tau fibrils formed from some of the most potent inducers: HS (**1**), polyphosphate (**13**), poly-L-glutamic acid (**22**), polyA (**24**), sodium alginate (**9**), and polystyrene sulfonate (**28**), were incubated for 60 min with the protease trypsin at a protein:protease ratio of 500:1 and then analyzed by SDS-PAGE. To distinguish fibril regions resistant to digestion, we probed these samples using several anti-tau antibodies (Tau 1, Tau 5, 4R, and Tau 13), which have epitopes that span the domains of tau (Fig 5a).

**Figure 5.**
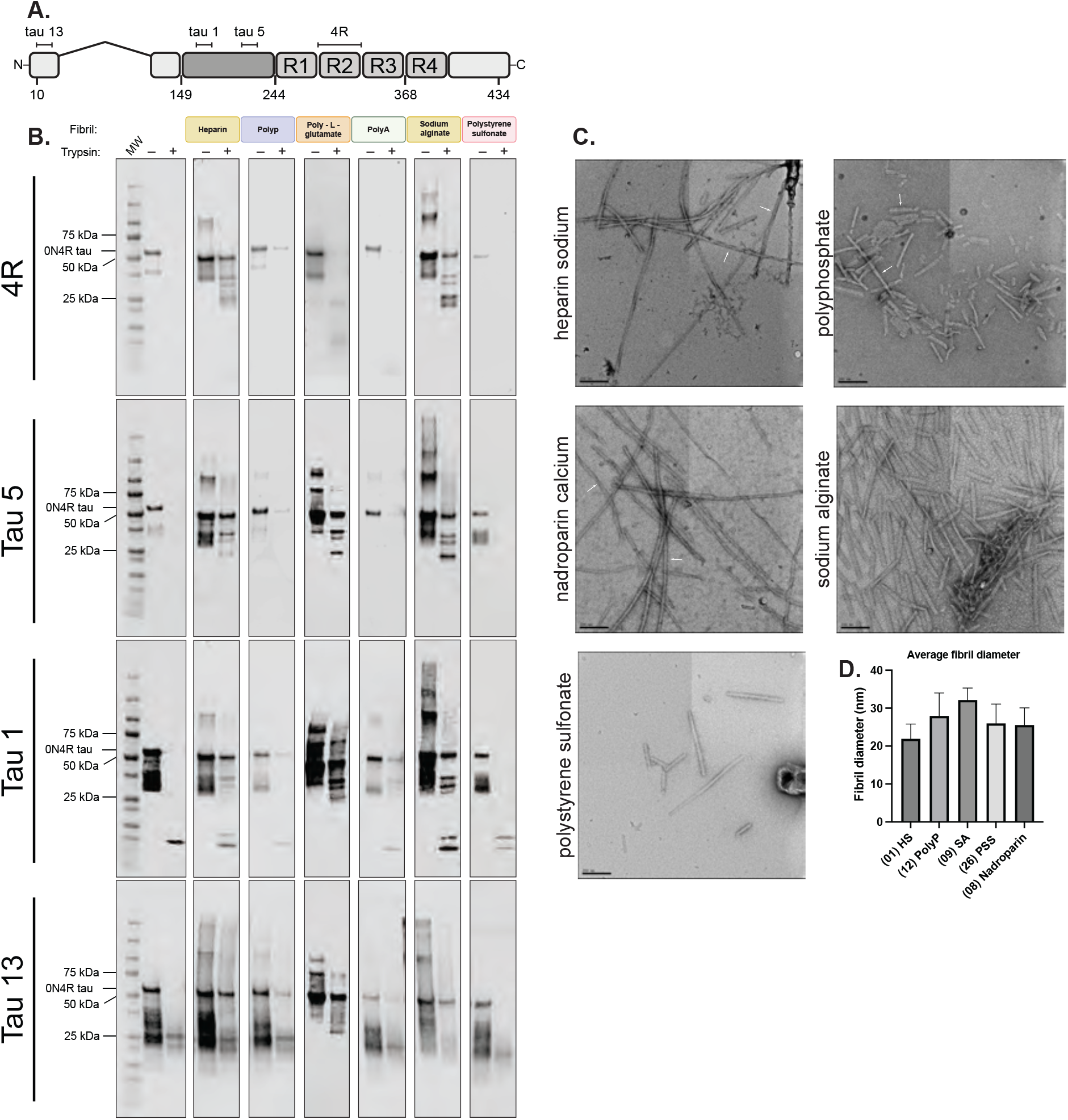
The identity of the polyanion impacts tau fibril structure. (A) The domain architecture of 0N4R tau, showing the location of the epitopes for anti-tau antibodies (anti-tau 13, 1, 5, and 4R). (B) Tau fibrils are generally resistant to proteolysis, but the digestion patterns depend on the identity of the inducer. The protease resistant fragmentation of 0N4R WT tau filaments differentially induced using heparin (01), polyphosphate (13), poly-L-glutamic acid (22), polyA (24), sodium alginate (09), or polystyrene sulfonate (28). Fibrils were prepared using the corresponding inducer at its EC50 value, purified by ultracentrifugation, and subsequently proteolyzed using trypsin. Proteolysis products were separated by SDS PAGE and probed using anti-tau antibodies. (C) Representative electron micrographs of negatively stained fibrils from recombinant 0N4R tau assembled in vitro at the end point of each reaction. Scale bar, 200 nm. (D) Quantification of the average diameter of recombinant fibrils (n = 30).

In the limited proteolysis experiments, we observed rapid degradation of soluble tau, which is consistent with its intrinsic disorder and many trypsin cleavage sites (Fig 5b; Supplemental Fig. 5). For the fibrils, we generally observed resistance to proteolysis and, more importantly, we observed differential proteolytic banding between samples. Conformational divergence is perhaps most apparent when probing the proline rich region (PRR). For example, using the anti-tau-5 antibody (residues 210-230) three bands ranging from 25 – 37 kDa are apparent for heparin- (**1**), poly-L-glutamic acid- (**22)** and sodium alginate- (**9**) induced samples, whereas no protease resistant banding is observed in this same region using polyphosphate (**13)**, polyA (**24)**, or polystyrene sulfonate (**28**). Probing further upstream in the PRR, using the anti-tau-1 antibody (residues 192 – 204), the initial similarities observed between **1** and **9**, and **22** diverge. Specifically, heparin (**1**) and sodium alginate (**9**) maintain a similar proteolysis profile with two bands between 30 to 37 kDa, in addition to the emergence of a ∼15 kDa band. For poly-L-glutamic acid (**22)**, however, the fibrils remain resistant to degradation in this region, with two intense bands appearing around 25 to 30 kDa. Interestingly, the 15 kDa fragment also appears in samples induced with polystyrene sulfonate (**28**) and this is the only region where this sample displayed protease resistance. Probing the N-terminal domain (NTD; residues 20 – 35) reveals that only the fibrils formed using poly-L-glutamic acid (**22**) were resistant to digestion; whereas all the rest were susceptible. This result suggests that the fibrils formed in the presence of poly-L-glutamic acid (**22**) have NTDs that are oriented in manner that shields them from proteolysis. Together, findings using these three antibodies (Tau 1, Tau 5 and Tau 13) support a conclusion in which inducers impact the conformation(s) of the fuzzy coat.

The R2 repeat region of tau is particularly important because previous work has shown that it is included in a subset of patient-derived core structures but excluded from others^12,44^. Interestingly, we observe that the R2 repeat is included in the protected core of fibrils formed in the presence of heparin (**1**) and sodium alginate (**9**), but that it is excluded from those formed using polyphosphate (**13**), poly-L-glutamic acid (**22**), polyA (**24**) or polystyrene sulfonate (**28**). Thus, it seems likely that only a subset of the inducers promote the formation of R2-containing fold. Indeed, our results with polyA (**24**) are consistent with cryo-EM evidence, which has shown that the R2 region is excluded from the cores of fibrils formed using RNA^45^.The biological and/or structural significance of R2 positioning is not yet known, but it is interesting that the identity of the polyanion can cause dramatic changes to R2 protease sensitivity.

Finally, we used negative stain transmission electron microscopy (EM) to examine and compare the supramolecular architecture of tau fibrils. These experiments also served the added goal of independently confirming whether the measured ThT signals were due to the formation of amyloid fibrils. We found that samples prepared using heparin (**1**), polyphosphate (**13**) and nadroparin calcium (**8**) had characteristic amyloid morphologies, including a mixture of twisted or flat fibrils (Fig 5c). The results with HS (**1**) are consistent with previous findings^46^. We also observed oligomeric (*i*.*e*. spherical) structures in samples formed from nadroparin calcium (**8**). In contrast, filaments generated using sodium alginate (**9**) and polystyrene sulfonate (**28**) are relatively monomorphic. Fibrils formed using sodium alginate (**9**) were particularly striking in their unusual and consistent morphology; moreover, they tended to produce thicker fibers (33 nm, n = 30) compared to the other inducers (Fig 5d). Together, these results support our hypothesis that polyanion identity strongly impacts the conformation of tau fibrils, such that the protein can be directed into strikingly different shapes by the chemical properties of the inducer.

## Discussion

The role(s) of anions in tau self-assembly have remained elusive. Many pioneering manuscripts have shown that individual polyanions promote this process^22,27,33,47,48^, with the most attention placed on heparins and polyphosphates. Here, we expand the landscape of tested anions, with a special focus on those naturally occurring ones that tau might encounter in the brain. We also tested them side-by-side to avoid interpretations that could arise due to differences in experimental conditions (*e*.*g*., buffer, tau concentration). From these screens, we found that tau aggregation is enhanced by a wide variety of anions. Indeed, the most pervasive theme is that anions with dramatically different scaffolds (*e*.*g*., sugar, polyphosphate, *etc*) are capable of supporting tau self-assembly. This finding suggests that degenerate physical features, such as valency and charge, are more important than the specifics of the scaffold from which the anions are displayed (*e*.*g*, polymer, micelle). For example, the sparsely charged hyaluronic acids (**3**-**6**; 1 charge per repeat) and the low valency compounds (**14, 29, 30**) were relatively poor at promoting fibril formation, but highly charged and multivalent molecules (**1, 2, 7, 13, 17, 19, 28**) from many chemical series can induce this process. Together, these results are broadly consistent with a model in which polyanion-assisted aggregation proceeds via minimizing the electrostatic repulsion between positively charged tau monomers, allowing subsequent inter- or intra-molecular scaffolding of monomers. Yet, the identity of the scaffold must also be important at some level because different anions produce distinct tau fibril morphologies. For scaffolds that have sufficient valency, it seems likely that more subtle features, such as flexibility or charge density, might then favor differential effects on lag time, elongation rate, and, ultimately, the structure of the resulting fibrils. For example, heparin sulfate (**1**) and sodium alginate (**9**) are both repeating di-saccharides, yet they produce dramatically different tau fibrils, as judged by either proteolysis or EM (see Fig 5). We speculate that the molecular details of the tau-anion contacts might be an important step in “folding” and fibril formation.

In the intact brain, soluble tau likely encounters many diverse and abundant biological molecules that bear a negative charge, including many of the proteins, lipids, nucleic acids, and metabolites tested here. Thus, it is interesting to consider the implications of the results in that context. For example, we find that the identity of the anion can have a strong impact on the structure of tau fibrils; thus, differences in the anion composition of specific brain regions could help dictate what types of fibrils are supported. In addition, it is possible that mixtures of anions, as likely encountered in complex cellular environments, could produce new structures or structures of mixed conformation. This is an important consideration because structural studies have shown that filaments formed *in vitro* using HS as an inducer do not have the same structure(s) as those isolated from patient brains.^46^ It is possible that different anions or combinations of anions might better replicate the patient-derived structures *in vitro*. Finally, we found that several polyanions, such as hyaluronic acids, are not capable of supporting aggregation at all. Similarly, low valency anions, such as triphosphate, were also remarkably poor at promoting tau aggregation. This is an important observation because these anions might serve as competitors *in vivo*; binding to cationic sites on tau and buffering the protein’s interactions with other, aggregation-promoting polyanions. If this supposition is true, then the relative concentrations of both active and inert anions might combine to dictate whether tau forms fibrils.

It is important to note that recombinant tau produced from *Escherichia coli* was used in these studies, so the proteins are devoid of post-translational modifications (PTMs). *In vivo*, the overall charge of tau will be tuned by multiple PTMs, including phosphorylation, acetylation, and ubiquitination^49,50^. These modifications can either add a negative charge (phosphate) or neutralize a positive one (acetylation), thus altering the isoelectric point and, in turn, adjusting tau’s sensitivity to polyanions^51^. Indeed, recombinant tau fibrils induced using HS (**1**) are significantly different than those formed using phosphorylated tau^52^. Future studies will be needed to deconvolute the impact of the vast number of possible PTM combinations. Likewise, mutations in tau might also impact its interactions with anions by altering the overall structural landscape of the protein. Indeed, we found that P301S tau was sensitive to a wider range of anions than WT, which might partly underlie this mutation’s important role in FTD. There are hundreds of additional mutations linked to tauopathy^53^, which might also vary in their response to polyanions.

More broadly, electrostatic forces are essential in mediating protein-protein interactions (PPIs)^54,55^, not just those found in tau. Here, we screened a library of naturally occurring and synthetic polyanions and found striking differences in how they promote tau self-assembly. It seems likely that a similar, screening-based approach could be adopted to probe the potential roles of polyanions on other PPIs. While crowding agents, such as polyethylene glycol, are sometimes explored^56^, systematic studies of polyanion libraries are less common.

## Supplementary Figure legends

**Supplementary Fig. 1** Identification of anions that produce Thioflavin T (ThT) artifacts. Each anion in the library was screened at a range of concentrations in the absence of tau protein to identify artifacts. (A) Endpoint RFUs were determined after 30 minutes at a single concentration. Several anions including **20, 21, 24, 25, 26**, and **29** generated ThT signal greater than 70 RFUs. This subset of anions was considered “ThT incompatible” and omitted from subsequent experiments. The one exception was chondroitin sulfate A (**02**), which we found to produce robust tau fibril formation. In this one case, the background ThT signal (∼447 RFUs) was subtracted. (B) In the absence of protein, the contribution of aberrant ThT fluorescence was confirmed over a minimum of 24 hours at several concentrations. Results are the average of technical replicates, and the error bars represent SEM (n = 3).

**Supplementary Fig. 2** A subset of molecules are unable to promote tau aggregation *in vitro*. The subset of anions that are unable to produce ThT positive species in the absence or presence of protein are deemed inert. (A) Under the tested conditions, hyaluronic acids of various molecular weights (**3-6**), glycolipids including ganglioside salts (**20 – 21**), multiple sugars (**11 – 12**), nucleic acids (**24 – 27**), fusidic acid (**29**), and bilirubin (**30**) do not generate aggregates of WT tau. (B) For mutant P301S tau, these same molecules do not initiate aggregation, excluding poly-L-lysine (**23**).

**Supplementary Fig. 3** A subset of anions weakly induce tau aggregation. (A) Using WT tau, fondaparinux (**7**), sodium alginate **(9)**, and diadenosine pentaphosphate (**15**) weakly stimulate fibril formation at 24 hours. Each anion produces a ?RFUs > 40, however, a complete sigmoidal curve was not reached. We categorize this behavior as “weak induction” (see text). (B) Similarly, diadenosine pentaphosphate (**15**) and poly-L-lysine (**23**) weakly stimulate aggregation of P301S tau.

**Supplementary Fig. 4** Several anions accelerate aggregation at low concentrations, but then inhibit it at higher concentrations (producing a “hook effect”). (A) For tauWT, a hook effect was observed using inducers **7 – 9, 21, and 28**. (B) Similar trends are identified using the tauP301S construct, however, for a larger subset of inducer molecules including **2, 7 – 10**, and **28**. Each curve represents are the average of three experiments performed in technical triplicate and the error bars represent SEM (n=3). As stated in the text, these inducers were used at EC50 values that were selected after excluding the higher concentrations.

**Supplementary Fig. 5** Full Western blots for the images shown in the partial proteolysis studies (Figure 5). The full, uncropped Western blots for the partial proteolysis studies, using antibodies (a) Tau 4R (b) Tau 5 (c) Tau 1 and (d) Tau 13. See text for details.

## Methods

### Recombinant protein expression and purification

The gene encoding human 0N4R tau was cloned into a pET-28a vector and transfected into *E. coli* BL21(DE3) competent cells. Starter cultures were grown in Luria broth (LB) containing 50 µg/mL kanamycin overnight at 37 °C with constant shaking at 200 rpm. Then, 20 mL of the starter culture was used to inoculate 1L of Terrific broth (TB), containing 50 µg/mL of kanamycin. Cells were grown at 37 °C, with constant shaking, until OD_600_ between 0.6 and 0.8 was reached. At this point, the incubation temperature was set to 30 °C, and NaCl (500 mM) and betaine (10 mM), were included in the growth medium. After 30 minutes, expression was induced with 200 µM IPTG for 3.5 h at 30 °C.

To purify tau, cells were pelleted and resuspended in a lysis buffer containing 20 mM MES (pH 6.8), 1 mM EGTA, 0.2 mM MgCl_2_, 5 mM DTT, and 1x cOmplete™ protease inhibitor cocktail (Roche). Cells were lysed by sonication and then boiled for 20 minutes. The lysate was clarified by centrifugation for 30 minutes at 30,000 x g. The clarified supernatant was dialyzed overnight into His binding buffer (1x PBS, 20 mM imidazole, 200 mM NaCl, and 5 mM β-mercaptoethanol (BME)) and purified by affinity purification.

His-tagged tau was bound to Ni-NTA resin for 1 h at 4 °C with constant mixing. The resin was washed using 500 mL of His binding buffer (1x PBS, 20 mM imidazole, 200 mM NaCl, and 5 mM BME), wash buffer 1 (1x PBS, 10 mM imidazole, 300 mM NaCl, and 5 mM BME) and, wash buffer 2 (1x dPBS, 15 mM imidazole, 100 mM NaCl, and 5 mM BME), and then eluted using 30 mL of elution buffer (1x PBS, 300 mM imidazole, 200 mM NaCl, and 5 mM BME). Tau was further purified using reverse-phase HPLC, as described previously.^14^ The protein was then lyophilized, resuspended in tau buffer (1x dPBS, 2 mM MgCl_2_, and 1 mM DTT). Protein concentration was determined using the bichinchronic acid (BCA) method.

### Compound Preparation

All compounds including anions and polyanions were sourced from commercial vendors and used without further purification. See Tables 1 and 2 for details of the catalog numbers. Each polyanion was freshly prepared in assay buffer (1x Dulbecco’s PBS pH 7.4, 2 mM MgCl_2_, 1 mM DTT) and sterilized with a 0.2-micron filter before each experiment. Lipid inducers (arachidonic acid, linoleic acid, phosphatidyl-L-serine) were handled similarly, except that they contained a final concentration of 5% ethanol to maintain solubility.

### Tau aggregation and kinetic screening

The ThT-based aggregation screen was performed in a miniaturized, 384-well plate format.^24^ The microplates (Corning 4511) were pre-rinsed with 20 µL 0.01% Triton-X to minimize interactions with the sides of the plate. In each well, tau (10 µM), thioflavin T (10 µM), polyanion (see Tables 1 and 2) for concentrations) and assay buffer (Dulbecco’s PBS pH 7.4, 2 mM MgCl_2_, 1 mM DTT) were added to each well, to a total volume of 20 µL. The aggregation reaction was carried out at 37 °C with continuous shaking and monitored via fluorescence (excitation=444 nm, emission=485 nm, cutoff=480 nm) in a Spectramax M5 microplate reader (Molecular Devices). Readings were taken every 5 min for at least 24 h. Each experiment was performed in triplicate wells. All components of the aggregation reaction were freshly prepared each day.

### Data processing

For data processing, the ThT signal produced by three replicates of the tau-only controls (no inducer) were averaged and this background was subtracted from corresponding samples. These values were typically less than 5 to 10% of the overall signal. For inducers with ThT background greater than 10%, mainly chondroitin sulfate A, inducer + ThT background was further subtracted. To identify the dose of inducer for subsequent kinetic analyses, we first determined the inducer concentration required to produce half-maximal ThT fluorescence signal by fitting the fluorescence curves to a sigmoid in GraphPad PRISM. Using the half maximal concentration, we analyzed the kinetic parameters of tau aggregation using the Grace plotting program (http://plasma-gate.weizmann.ac.il/Grace/) and fitting the aggregation curves to the Gompertz function: y=Ae^(-e^((t-ti_i)/b))^, as previously described^24^. In that equation, the lag-time is defined by the inflection point; the inverse of apparent elongation rate constant (ti-b). In addition, A represents the maximum ThT signal, and the apparent elongation rate constant is 1/b. For error analysis, we considered the kinetic measurements from individual experiments and used them to calculate standard error of the mean (SEM), without considering additional error introduced by goodness of fit.

### Fibril preparation for proteolysis

The aggregation reaction was performed in 1.5 mL Eppendorf tubes for 48 hours with constant agitation at 1200 rpm. The reactions included tau (10 µM) and inducer (see Tables 1 and 2 for concentrations) in assay buffer (Dulbecco’s PBS pH 7.4, 2 mM MgCl_2_, 1 mM DTT) with a total volume of 300 µL. After 48 hours, the reactions were subjected to ultracentrifugation (Beckman Optima™ Max-XP Tabletop Ultracentrifuge) using a TLA-55 Fixed-Angle Rotor at 103,000 rcf to remove monomeric tau and excess inducer. Pelleted tau fibrils were then resuspended in 1/3-1/4 of the reaction volume and the concentration was determined by Coomassie band intensity measured against a standard and quantified in ImageLab (BioRad).

### Limited proteolysis

Soluble tau and freshly prepared fibrils were digested with Promega sequencing grade trypsin at a protein:protease ratio 500:1 for 60 minutes at 37 °C. The reaction was performed in 40 mM HEPES 40 mM NaCl pH 8.0. Reactions were quenched with 3X SDS loading buffer + 1 mM PMSF and immediately heat denatured at 95 °C for 5 minutes. The proteolysis reactions were separated using a 4-20% polyacrylamide Tris-glycine SDS-PAGE denaturing gel (Invitrogen). Gels were transferred to nitrocellulose using a TurboBlot (BioRad) and analyzed using antibodies corresponding to several tau epitopes (anti-tau 1, 5, 13 and anti-tau 4R). Mouse anti-tau 1, 5 (Thermo) and rabbit anti-4R tau (Abcam) were prepared 1:1000 in Intercept T20 (TBS) Antibody Diluent (LiCor) and anti-tau 13 (Abcam) was prepared 1:5000. All secondary antibodies were prepared 1:10000 in 1:1 TBST and T20 (TBS) Antibody Diluent (LiCor).

### Transmission electron microscopy

0N4R tau^WT^ fibrils were freshly prepared as described above in the kinetic screening assay, for 36 hours without ThT present. The corresponding samples were pooled and subsequently immobilized on 600-mesh carbon-coated copper grids (SPI). The samples were incubated for 30 seconds on a glow-discharged grid, and then the solution was removed by filter paper. Three washing steps with double distilled H_2_O were followed by three staining steps with 0.75% (w/v) uranyl formate (Electron Microscopy Sciences). The samples were imaged using a FEI Tecnai 10 operated at 100 keV. Micrograph images were recorded using a 4kx 4k CCD camera (Gatan). The fibril dimensions were measured using the ImageJ software.

## Author Contributions

Kelly Montgomery: experimental design, reagent preparation, perform experiments, interpret results, manuscript draft, manuscript preparation

Emma Carroll: experimental design, reagent preparation, perform experiments, interpret data, manuscript preparation

Aye Thwin: perform experiments, interpret data

Paige Hodges: perform experiments, interpret data

Daniel Southworth: manuscript preparation

Jason Gestwicki: experimental design, interpret data, manuscript preparation

## Acknowledgements

The authors thank Daniel Schwarz, Taia Wu, Julia Jones, and Jason Hernandez for technical assistance. We also acknowledge funding from the Tau Consortium (to J.E.G. and D.R.S.), the HHMI Gilliam predoctoral Fellowship and Ford Foundation Predoctoral fellowship (to K.M.M.), and the National Institute on Aging NRSA F32 (to E.C.C, F32AG076281).

## COI

The authors declare no conflict of interest

## Notes

### Competing Interest Statement

The authors have declared no competing interest.

