## Supporting Information for "The Chemical Features of Polyanions Modulate Tau Aggregation and Conformational States"

**Supplementary Fig. 1** Identification of anions that produce Thioflavin T (ThT) artifacts.

**Supplementary Fig. 2** A subset of molecules are unable to promote  tau aggregation *in vitro*.

**Supplementary Fig. 3** A subset of anions weakly induce tau aggregation.

**Supplementary Fig. 4** Several anions accelerate aggregation at low concentrations, but then inhibit it at higher concentrations (producing a “hook effect”).

**Supplementary Fig 5.** Full Western blots for the images shown in the partial proteolysis studies.

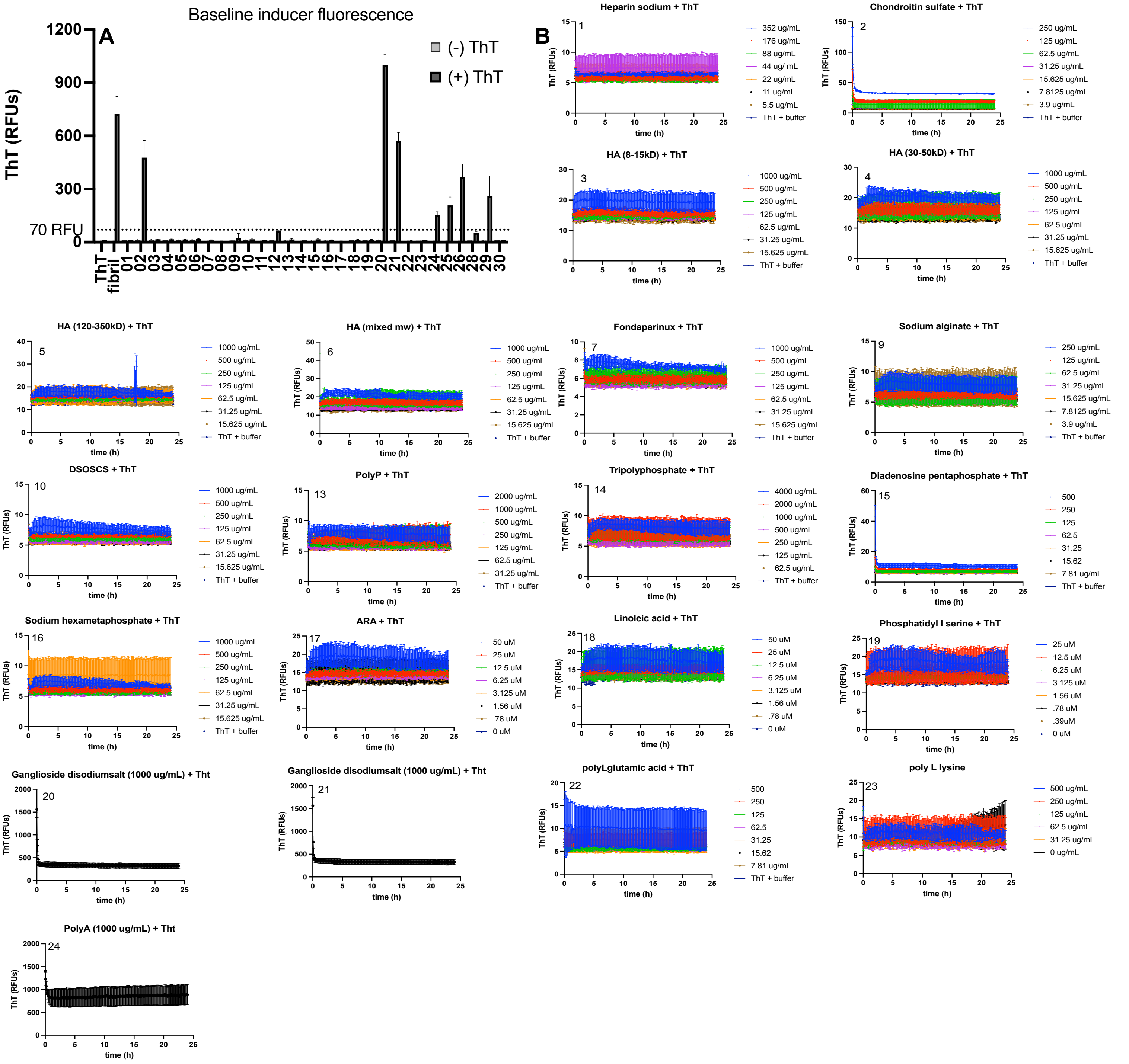

**Supplementary Fig. 1** Identification of anions that produce Thioflavin T (ThT) artifacts. Each anion in the library was screened at a range of concentrations in the absence of tau protein to identify artifacts. (A) Endpoint RFUs were determined after 30 minutes at a single concentration. Several anions including 20, 21, 24, 25, 26, and 29 generated ThT signal greater than 70 RFUs. This subset of anions was considered "ThT incompatible" and omitted from subsequent experiments. The one exception was chondroitin sulfate A (02), which we found to produce robust tau fibril formation. In this one case, the background ThT signal (~447 RFUs) was subtracted. (B) In the absence of protein, the contribution of aberrant ThT fluorescence was confirmed over a minimum of 24 hours at several concentrations. Results are the average of technical replicates, and the error bars represent SEM (n = 3).

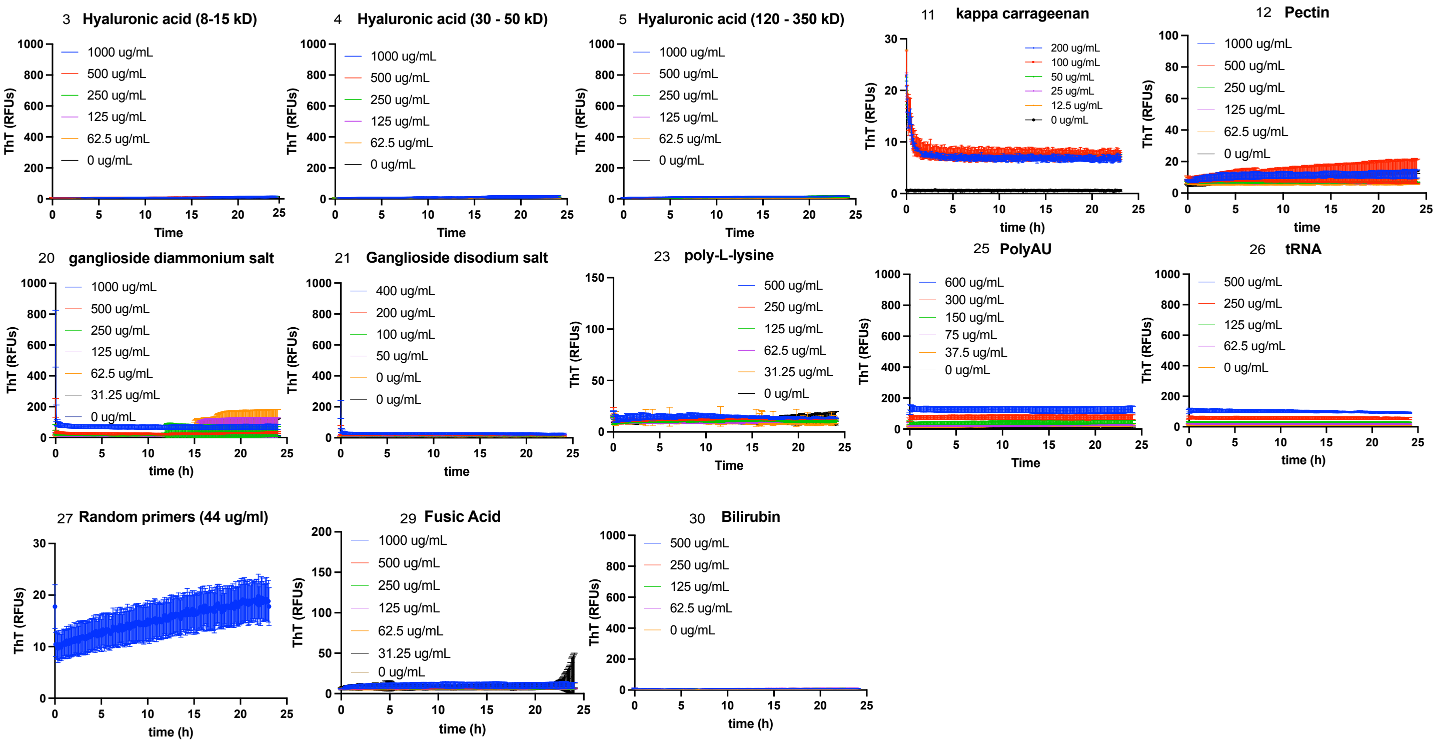


**Supplementary Fig. 2** A subset of molecules are unable to promote tau aggregation *in vitro*. The subset of anions that are unable to produce ThT positive species in the absence or presence of protein are deemed inert. (A) Under the tested conditions, hyaluronic acids of various molecular weights (3-6), glycolipids including ganglioside salts (20 – 21), multiple sugars (11 – 12), nucleic acids (24 – 27), fusidic acid (29), and bilirubin (30) do not generate aggregates of WT tau. (B) For mutant P301S tau, these same molecules do not initiate aggregation, excluding poly-L-lysine (23).

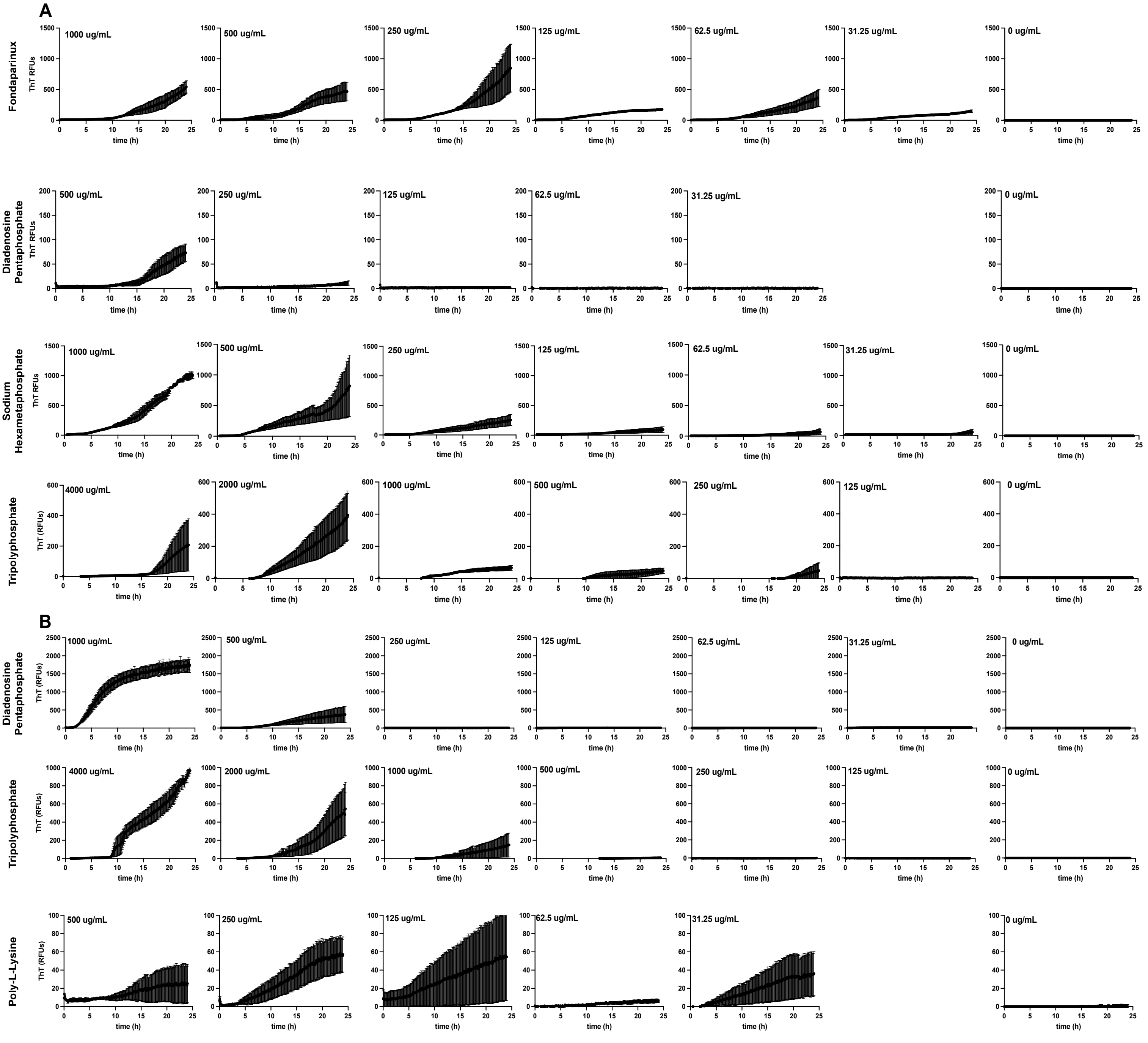


**Supplementary Fig. 3** A subset of anions weakly induce tau aggregation. (A) Using WT tau, fondaparinux (7), sodium alginate (9), and diadenosine pentaphosphate (15) weakly stimulate fibril formation at 24 hours. Each anion produces a ΔRFUs > 40, however, a complete sigmoidal curve was not reached. We categorize this behavior as “weak induction” (see text). (B) Similarly, diadenosine pentaphosphate (15) and poly-L-lysine (23) weakly stimulate aggregation of P301S tau.

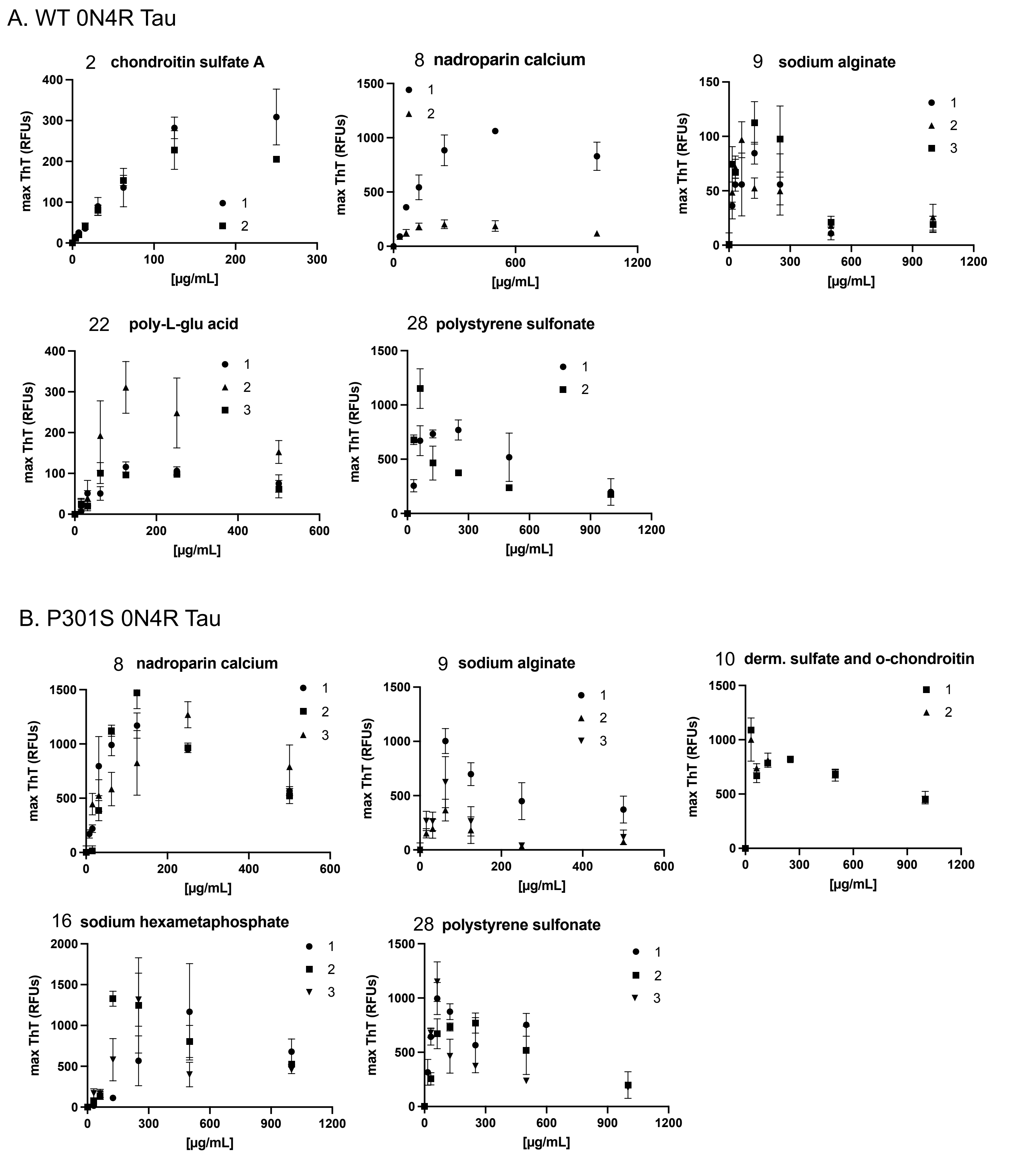


**Supplementary Fig. 4.** Several anions accelerate aggregation at low concentrations, but then inhibit it at higher concentrations (producing a “hook effect”). (A) For tauWT, a hook effect was observed using inducers 7 – 9, 21, and 28. (B) Similar trends are identified using the tauP301S construct, however, for a larger subset of inducer molecules including 2, 7 – 10, and 28. Each curve represents are the average of three experiments performed i technical triplicate and the error bars represent SEM (n=3). As stated in the text, these inducers were used at EC50 values that were selected after excluding the higher concentrations.

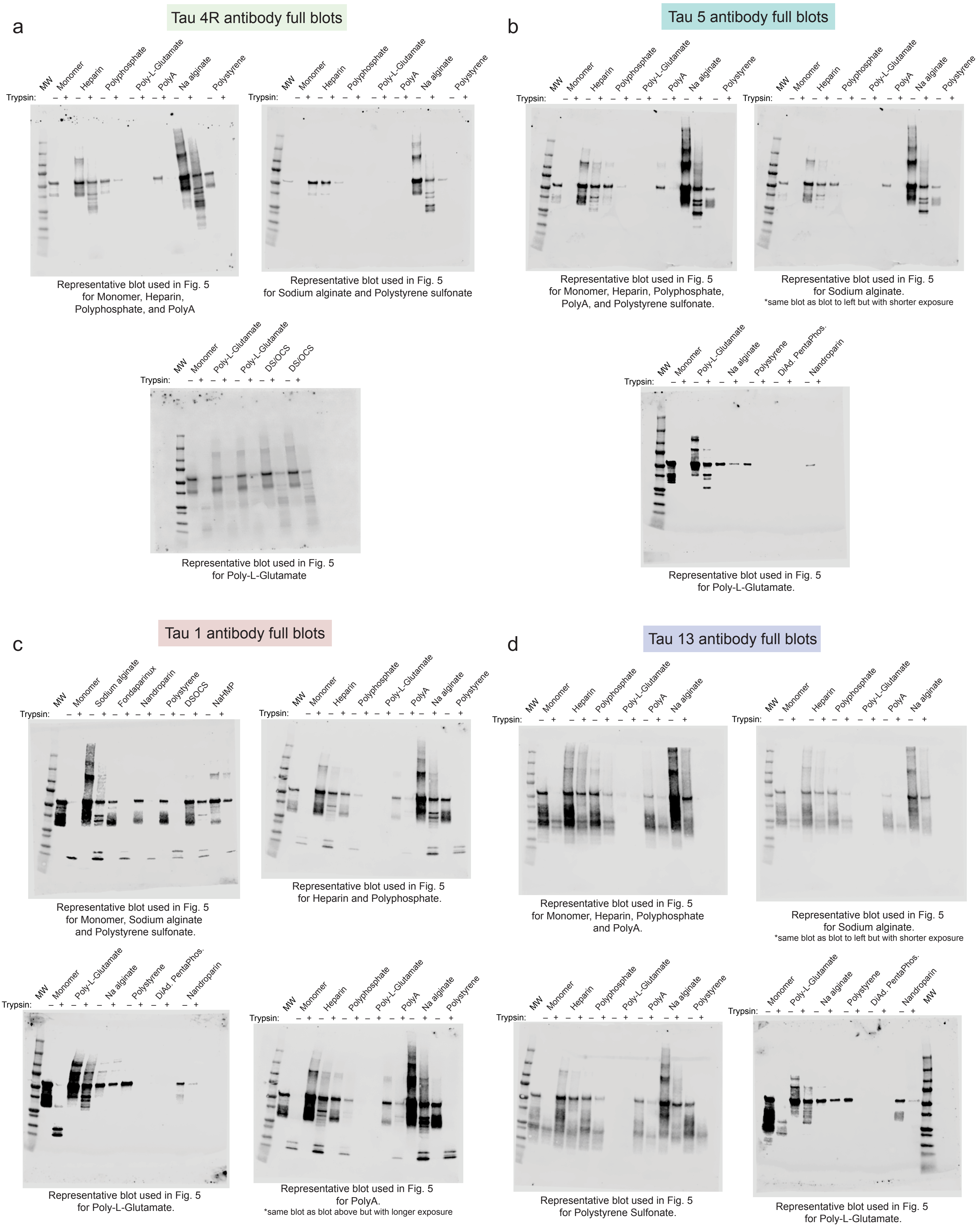


**Supplementary Fig. 5** Full Western blots for the images shown in the partial proteolysis studies (Figure 5). The full, uncropped Western blots for the partial proteolysis studies, using antibodies (a) Tau 4R (b) Tau 5 (c) Tau 1 and (d) Tau 13. See text for details.
